# Understanding how genetically encoded tags affect phase separation by Heterochromatin Protein HP1α

**DOI:** 10.1101/2023.12.04.569983

**Authors:** Ziling (Kate) Zhou, Geeta J. Narlikar

## Abstract

Liquid-liquid phase separation (LLPS) is driven by weak multi-valent interactions. Such interactions can result in the formation of puncta in cells and droplets *in vitro*. The heterochromatin protein HP1α forms droplets with chromatin *in vitro* and is found in puncta in cells. A common approach to visualize the dynamics of HP1α in cells is to genetically encode fluorescent tags on the protein. HP1α modified with tags such as GFP has been shown to localize to dynamic puncta *in vivo*. However, whether tagged HP1α retains its intrinsic phase separation properties has not been systematically studied. Here, using different C-terminal tags (AID-sfGFP, mEGFP, and UnaG), we assessed how tag size and linker length affected the phase separation ability of HP1α with DNA *in vitro*. We found that the AID-sfGFP tag (52 kDa) promoted HP1α phase-separation, possibly driven by the highly disordered AID degron. The mEGFP tag (27 kDa) inhibited phase-separation by HP1α, whereas an UnaG tag (13 kDa) with a 16 amino acid linker showed minimal perturbation. The UnaG tag can thus be used in cellular studies of HP1α to better correlate *in vitro* and *in vivo* studies. To test if cellular crowding overcomes the negative effects of large tags *in vivo*, we used polyethylene glycol (PEG) to mimic crowding *in vitro*. We found that addition of 10% PEG8000 or PEG4000 enables phase separation by GFP-tagged HP1α at comparable concentrations as untagged HP1α. However, these crowding agents also substantially dampened the differences in phase-separation between wild-type and mutant HP1α proteins. PEG further drove phase-separation of Maltose Binding Protein (MBP), a tag often used to solubilize other proteins. These results suggest that phase-separation of biological macromolecules with PEG should be interpreted with caution as PEG-based crowding agents may change the types of interactions made within the phases.

## Introduction

The cell is a crowded space consisting of different intracellular organelles that are involved in distinct physiological processes (1). The cell utilizes membrane-bound organelles such as the nucleus to temporally and spatially separate biochemical reactions. There are also other cellular compartments that are not membrane-bound such as P-bodies and nucleoli in the cytoplasm and nucleus, respectively (2, 3). Such membrane-less organelles are thought to form through the process of liquid-liquid phase separation (LLPS) which results in puncta formation in cells and droplets *in vitro* (4). LLPS is driven by weak multi-valent interactions between macromolecules. Many factors play important roles in promotion of LLPS, including the modularity of protein domains, polymer multivalency and the presence of intrinsically disordered regions (IDRs) (5).

Chromatin compartmentalization via LLPS in the nucleus has been implicated in regulating gene expression (6). Many proteins involved in this process have an intrinsic ability to form droplets. The mammalian heterochromatin protein, heterochromatin protein 1 (HP1α), is involved in gene silencing by compacting and spatially segregating heterochromatin (7-10). *In vitro*, HP1α can undergo LLPS when its N-terminus is phosphorylated or when it interacts with chromatin or DNA. However, the roles of the LLPS properties of HP1α in heterochromatin structure and function *in vivo* are not well-understood. Visualizing HP1α dynamics in cells is important for understanding how HP1α mediated chromatin droplets *in vitro* relate to *in vivo* function. To enable such visualization, prior studies have genetically encoded fluorescent tags on HP1α in cells (7, 10-13). The phase-separation of HP1α depends on multivalent interactions, between HP1α molecules, which could be influenced by the introduction of a large tag. Indeed, it has been shown that the addition of a GFP tag to either the N or C-terminus of phosphorylated HP1α inhibits its ability to phase separate *in vitro* (8). This raises the question of whether tagged HP1α retains its native functions and interactions in a cell. It is therefore crucial to identify a tagging arrangement on HP1α that minimally perturbs its biophysical properties *in vitro* as this will allow for a more rigorous interpretation of *in vivo* results.

It is currently unclear whether the inhibitory effects of the GFP tag on HP1α reflect a general consequence of adding a tag, or whether there are alternative arrangements of tags that can permit phase separation at concentrations similar to untagged HP1α (8). To investigate this possibility, we have systematically examined the effects of different C-terminal fluorescent tags (AID-sfGFP, mEGFP, and UnaG) on HP1α’s phase separation ability *in vitro* (Figure 1) (14, 15). These tags vary in size (13 – 52 kDa) and we further varied the length of the linker (8 – 16 aa) used to separate the tag from HP1*α*.

**Figure 1.**
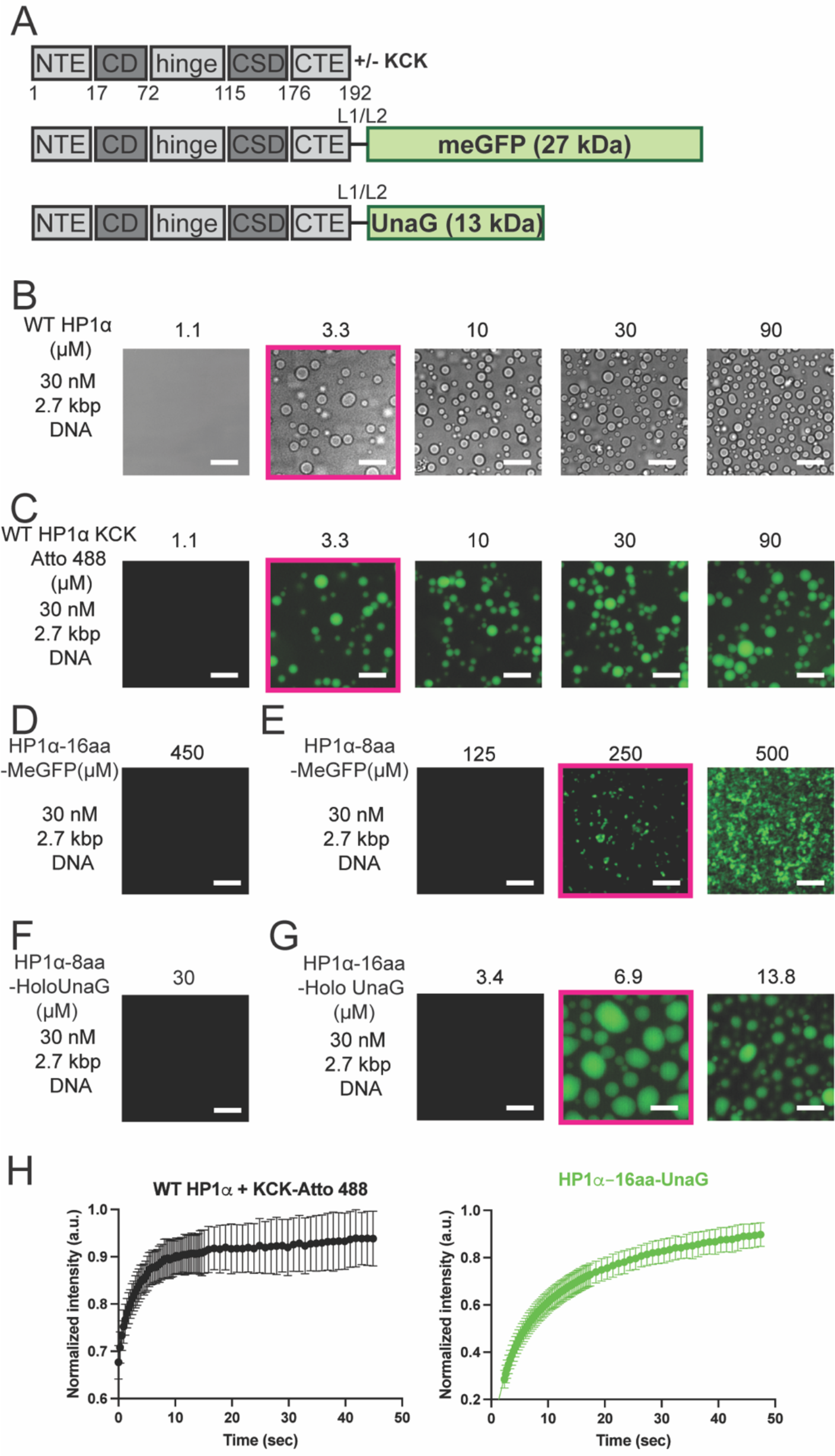
Tag size and linker length influence HP1α’s phase separation properties *in vitro*. (A) Domain diagrams of WT HP1α with or without tags. L1 and L2 represent linker lengths of 8 and 16 amino acids, respectively. Sequence of the linkers are GSGGSGGS and GSGGSGGSGSGGSGGS. (B) Bright field images of a concentration series of WT HP1α with 30 nM 2.7 kbp DNA. The red box indicates the saturation concentration for phase separation. (C) Fluorescent images of a concentration series of HP1α-KCK-Atto488 (1 WT:50 KCK) with 2.7 kbp DNA. (D) Fluorescent image of 450 *µ*M HP1α-16aa-mEGFP with 2.7 kbp DNA. (E) HP1α-8aa-mEGFP with 2.7 kbp DNA. (F) Fluorescent image of HP1α-8aa-(holo)UnaG mixed with 2.7 kbp DNA. (G) Fluorescent images of serial dilution of HP1α-16aa-(holo)UnaG with 2.7 kbp DNA. (H) FRAP analysis of WT HP1α (left) and HP1α-16aa- (holo)UnaG (right) condensates. N of 8. Scale bar represents 20 *µ*m.

Within our series of tags, the smallest fluorescent tag, UnaG (13 kDa), with a 16 amino acid (aa) linker was the only tag that minimally perturbed HP1α’s phase separation ability *in vitro*. However, larger tags that inhibit *in vitro* phase separation can still form puncta in a cell (11, 12). We reasoned that this discrepancy could be explained by cellular crowding. To recapitulate cellular crowding, we introduced a commonly used crowding agent polyethylene glycol (PEG) (16-18). We found that PEG induces phase separation of HP1α constructs that have no intrinsic propensity to phase separate. Overall, our findings highlight the need to quantitatively test the phase-separation effects of fluorescent tags used to visualize HP1α in cells and raise questions about the biological relevance of condensates induced with PEG.

## RESULTS

### Effects of tag size and linker length on phase-separation of HP1*α*

HP1α has a domain architecture as shown in Figure 1A. The protein has two structured domains, the chromoshadow domain (CSD) and the chromodomain (CD). The CSD is responsible for homodimerization, and the CD recognizes the H3K9me3 mark on chromatin. The CSD and CD are connected by an intrinsically disordered region (IDR), termed the hinge domain that binds to nucleic acids. Two additional IDRs are present on the N-and-C termini of the protein, and are termed the N-terminal extension (NTE) and the C-terminal extension (CTE), respectively. Previous work has suggested that HP1*α* phase separates through interactions between the NTE and the hinge (8, 19). Additionally, the CTE interacts with the hinge domain in the absence of DNA to play an autoinhibitory role of inhibiting oligomerization (8). Truncation of the NTE substantially inhibits HP1α’s phase separation while deletion of the CTE promotes HP1α’s phase-separation (19). The need for an intact NTE to allow phase-separation suggested that a tag would be better tolerated on the CTE. Therefore, all tags for this study were placed on the C-terminus of HP1α.

Of the many variables related to a tag that could affect the ability of HP1α to phase separate, we focused on two, the size of the tag and the length of the linker. We reasoned that varying these two features would vary the extent of steric hindrance in the dynamic higher order interactions required to drive phase separation. To address the effect of tag size, we chose two tags, monomeric EGFP (mEGFP, 27 kDa) and UnaG (13 kDa). We chose mEGFP to avoid the complication of GFP dimerization, as that could have separate effects on phase separation. To address the linker length effect, we tested linker lengths of 8 and 16 amino acids. We uncoupled the effects of tag size and linker length by designing four tagged HP1α constructs (HP1α-8aa-mEGFP, HP1α-16aa-mEGFP, HP1α-8aa-UnaG, and HP1α-16aa-UnaG) (Figure 1A).

We performed phase separation assays in the presence of a 2.7 Kbp plasmid DNA using a defined series of HP1α concentrations (19). Within this series, we define the saturation concentration for phase separation as the lowest concentration of HP1*α* at which we can detect droplet formation via microscopy. We compared phase-separation in the following buffer conditions: 20 mM HEPES, 70 mM KCl, 1 mM DTT. As previously reported, unlabeled HP1α readily undergoes phase separation with the 2.7 Kbp DNA construct, phase separating at a saturation concentration of 3.3 *µ*M (Figure 2B) (19). In addition to untagged HP1α, we also chemically labeled HP1α. We attached a KCK tag to the C-terminus of HP1α and labeled the cysteine residue with Atto488 using maleimide chemistry. HP1α-KCK-Atto488 also displayed a saturation concentration of 3.3 *µ* M, indicating that the chemically attached label does not perturb the phase separation properties of HP1α (Figure 2C). In contrast, HP1α-16aa-mEGFP remains soluble at high micromolar concentrations (450 *µ*M) (Figure 2D). With a shorter linker of 8aa, HP1α-8aa-mEGFP, forms non-spherical shaped phases at a saturation concentration of 250 *µ*M (Figure 2E). This is ∼75 fold higher than the saturation concentration of WT HP1*α*. These results indicate that a C-terminal mEGFP tag substantially interferes with the phase separation behavior of HP1α, independent of linker length. To test if the mEGFP tag affects binding of HP1α to DNA, we measured binding to a 187 bp stretch of DNA by electrophoretic mobility shift assays (EMSA) (Figure S1C and D). The K_1/2_ values for untagged HP1α and HP1α-16aa-mEGFP are 0.88 *µ*M and 0.58 *µ*M, respectively (Figure S1 C and D). The similar K_1/2_ values suggest that the mEGFP tag does not interfere with DNA binding but may affect oligomerization of HP1α.

**Figure 2.**
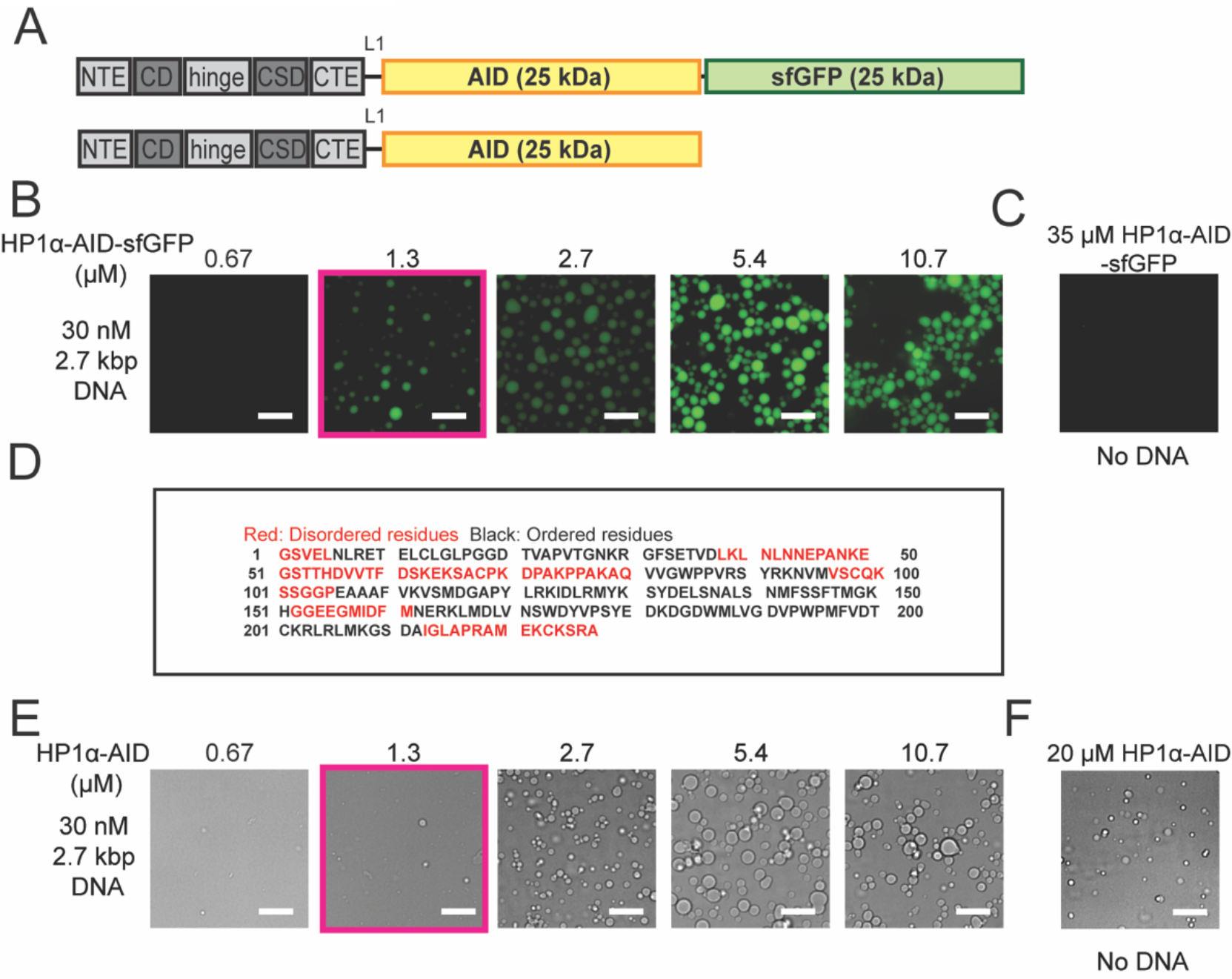
Large tags affect phase separation by HP1α. (A) Domain diagrams of HP1α with 2 different large tags, where L1 8 amino acids, respectively. (B) Fluorescent microscopy images of serial dilution HP1α-AID-sfGFP with 30 nM 2.7 kbp DNA. Red box indicates saturation concentration for phase separation. (C) Fluorescent image of 20 *µ*M HP1α-AID-sfGFP without DNA. (D) Sequence of the AID and disorder prediction from alpha-fold, where red colored letters are predicted to be highly disordered and black are predicted to be ordered. (E) 20X Brightfield images of a concentration series of HP1α-AID with 30 nM 2.7 kbp DNA. (F) Brightfield microscopy of 20 *µ*M HP1α-AID without DNA. Scale bar = 20 *µ*m.

Next, we evaluated the effects of the smaller UnaG tag. The UnaG protein shows fluorescent properties upon binding a cofactor, Bilirubin (Br)(15). Therefore, we tested the effects of the UnaG tag with (holo) and without (apo) its cofactor Br and connected via either an 8 aa or 16 aa linker. Without Br, HP1α-16aa-(apo)UnaG and HP1α-8aa- (apo)UnaG phase separate with saturation concentrations of 6.5 and 4.7 *µ*M, respectively (Figure S1 A and B). These values are comparable to the saturation concentrations of wild type HP1α, suggesting that the apo form of UnaG minimally perturbs HP1α’s phase separation function. However, in the holo form, HP1α-8aa-(holo)UnaG does not phase separate up to concentrations as high as 30 *µ*M. In contrast, HP1α-16aa-(holo)UnaG shows a similar saturation concentration (6.5 *µ*M) as the apo form (Figure 1F). Thus, in the series tested above, the least perturbative fluorescent tagging arrangement for HP1α is HP1α-16aa-(holo)UnaG.

We next assessed whether the UnaG tag affected the dynamics of HP1α within the phase separated droplets. We used fluorescence recovery after photobleaching (FRAP) on pre-formed phase separated HP1α-16aa-(holo)UnaG and HP1α-KCK-Atto488 droplets in the presence of DNA. The half-times for recovery of fluorescence in the fast phase are 1.9 s and 3.1 s, for HP1α-KCK-Atto488 and HP1α-16aa-UnaG, respectively. The two constructs also show comparable half-times for recovery of the slow phases (Table 3). These results indicate that this UnaG tag arrangement maintains the seconds time-scale dynamics of HP1α in its phase separated droplets and suggest that both size of the tag and linker length affect HP1α’s biophysical properties. Importantly, our studies have identified a specific arrangement of the UnaG tag that minimally perturbs phase separation by HP1α. This tag now allows direct comparison of the behavior of HP1α inside and outside of a cell.

More recently GFP tags have been combined with degrons to study effects of rapid protein degradation (14, 20). In the context of HP1α the AID-sfGFP tag (where AID is “Auxin Inducible Degron”), has been used to study the effects of rapidly degrading HP1ɑ in cells (Figure 2A). In HP1α-AID-sfGFP, AID and sfGFP combined are 50 kDa, around twice the size of HP1α. We therefore hypothesized that similar to mEGFP, this large tag would interfere with HP1α’s phase separation ability. Surprisingly, HP1α-AID-sfGFP forms phase-separated droplets in the presence of DNA at a lower saturation concentration (1.3 *µ*M) than untagged HP1α (Figure 2B and C). We hypothesized that the AID-sfGFP tag promotes phase separation due to a large intrinsic disordered region contributed by the AID (Figure 2D). To separately test AID’s effect on HP1α phase separation, we generated an HP1α-AID construct (Figure 2A). Unlike untagged HP1α, HP1α-AID now phase separates in the absence of DNA (Figure 2F). Further, HP1α-AID shows the same saturation concentration as HP1α-AID-sfGFP in the presence of DNA (Figure 2E). These results indicate that the AID tag enhances HP1α’s intrinsic phase separation ability and potentially rescues some of the inhibition introduced by the GFP tag in cells.

### Testing the effect of PEG8000 on phase separation properties

Interestingly, even though previous studies and our studies here show that a GFP tag inhibits phase-separation by HP1α, GFP-tagged HP1α constructs have been shown to be components of dynamic heterochromatin puncta *in vivo* (11, 13, 21-23). It is possible that phase separation of these tagged HP1α proteins in cells is promoted by other cellular contents and by the high degree of molecular crowding. To mimic cellular crowding *in vitro*, we used polyethylene glycol (PEG), a commonly used water-soluble macromolecular crowing agent. We tested two different PEG molecules (PEG·8000 and PEG·4000).

We first determined if PEG8000 promotes phase separation of HP1α without DNA. We titrated PEG and found that 10% is sufficient for HP1α phase separation alone (Figure 3A). This percentage of PEG is similar to that used in previous studies (16, 17, 24-26). In the absence of PEG, HP1α alone does not phase separate at concentrations up to 800 *µ*M (8, 19). However, adding 10% PEG8000 lowered HP1α’s saturation concentration to 3.3 *µ* M (Figure 3B). Analogously, PEG also enables HP1α-16aa-mEGFP to phase separate at 3.3 *µ*M without DNA (Figure 3B). To find a negative control, we used an HP1α protein with a chromoshadow mutation (CSDm). This mutation inhibits HP1α dimerization and the HP1α CSDm protein does not phase separate with the 2.7kb DNA construct, even at 147 *µ*M (Figure 3C) (8). However, PEG induces phase separation of HP1α CSDm in the absence of DNA and at a saturation concentration of 22.5 *µ*M, which is ∼6.8 fold higher than the other HP1*α* constructs above (Figure 3D).

**Figure 3.**
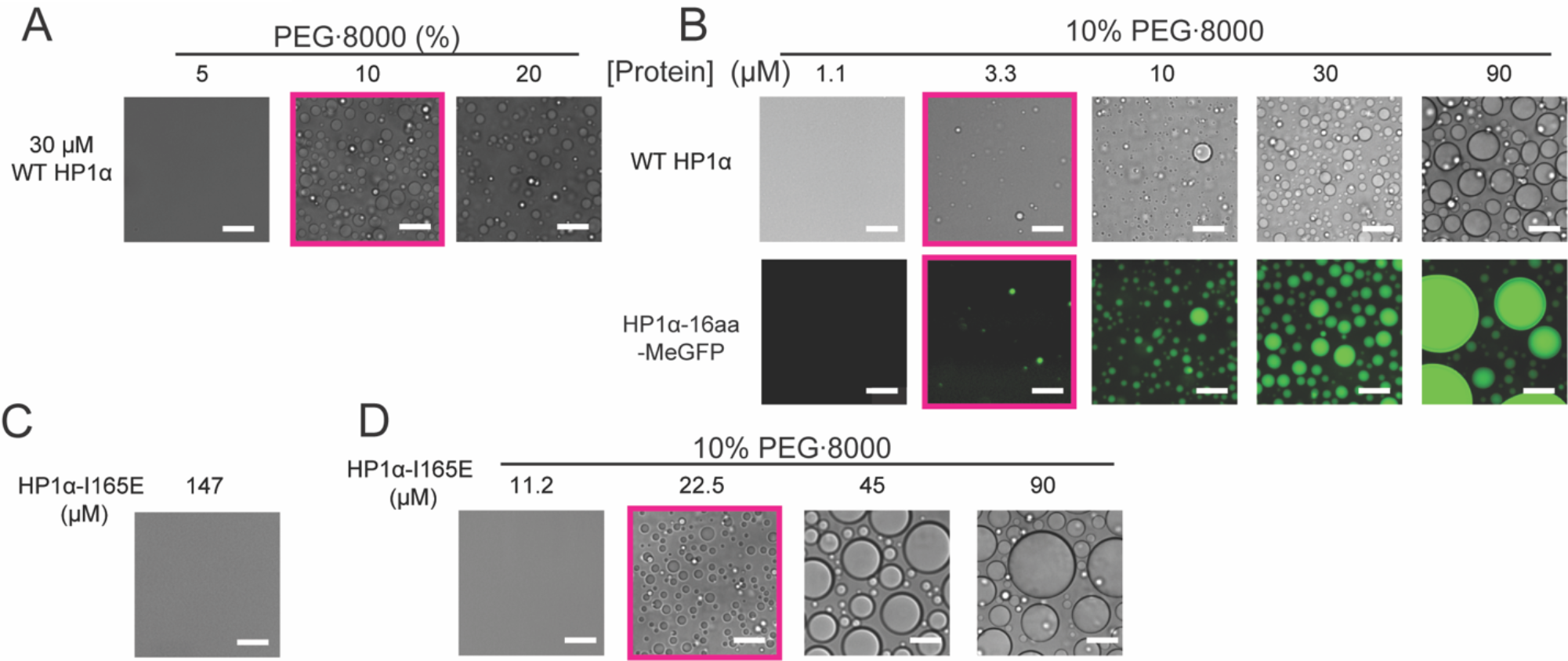
PEG8000 induce phase separation of HP1α and tagged constructs in the absence of DNA. (A) Three different PEG8000 percentages used to determine the minimum PEG percentage (10%) needed to induce HP1α phase separation without DNA. (B) Serial dilution of wt HP1α and HP1α-16aa-mEGFP with 10% PEG 8000. All three constructs now have similar saturation concentrations within a 3-fold difference. Red box represents saturation concentration for phase separation. (C) Bright field image of HP1α CSDm alone at 147 *µ*M with no visible droplets. (D) Bright field images of a concentration series of HP1α CSDm with 10% PEG8000, HP1α CSDm now displays phase separation starting at 22.5 *µ*M. Scale bar = 20 *µ*m.

Since PEG is also known to precipitate DNA, we investigated how it affects the phase-separation of HP1α in the presence of DNA. Brightfield microscopy revealed that under the conditions of our experiments PEG·8000 does not promote phase separation of DNA alone (Figure 4B). In the presence of PEG, DNA does not further lower the saturation concentration of untagged HP1α (Figure 4A). For HP1α-16aa-mEGFP, addition of DNA also did not decrease the saturation concentration in comparison to PEG and protein alone (Figure 4A). These results suggest that PEG8000 alters the protein-protein and protein-nucleic acid interactions within the droplets. We next studied the effect of PEG8000 on the HP1α CSDm with DNA since PEG induced phase separation of HP1α CSDm alone. We found that in the presence of PEG8000 HP1α CSDm now phase separates at 5.6 *µ*M, an ∼ 4-fold lower concentration than in the absence of DNA (Figure 4D).

**Figure 4.**
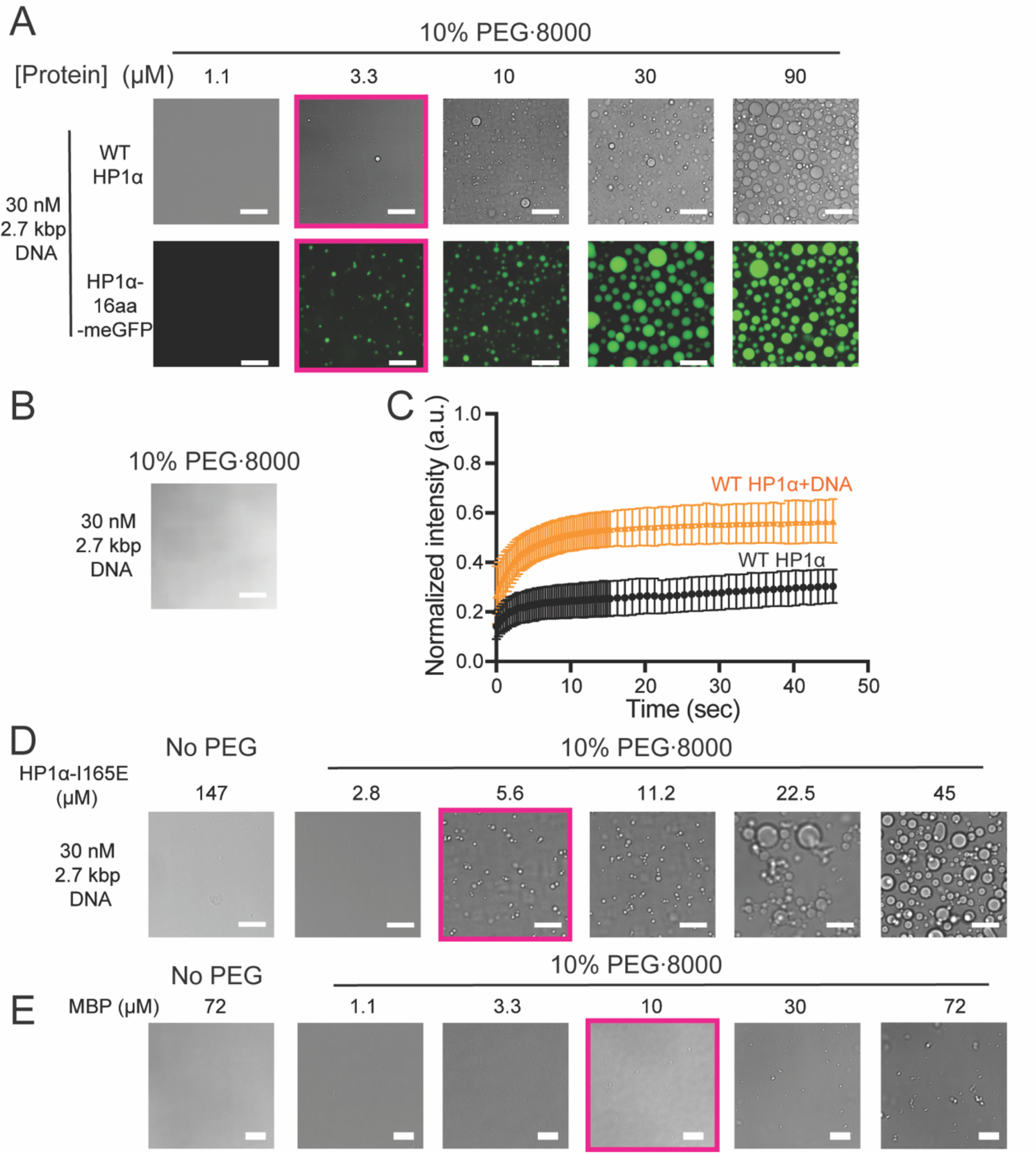
PEG8000 diminishes effects of DNA in promoting phase separation. (A) Serial dilution of WT HP1α with 30 nM 2.7 kbp DNA and 10% PEG8000. Scale bar = 20 *µ*m. Red box indicates saturation concentration for phase separation. (B) 10% PEG8000 with 30 nM 2.7 kbp DNA. (C) FRAP of PEG8000-induced HP1α-KCK-Atto488 and HP1α-KCK-Atto488 plus DNA droplets, n = 8. (D) Left, bright field image of 147 *µ*M HP1α CSDm with 30 nM 2.7 kbp DNA with no PEG 8000. Right, a concentration series of HP1α CSDm with 30 nM 2.7 kbp DNA with 10% PEG8000. Scale bar = 20 *µ*m. (E) Left, Maltose binding protein alone at 72 *µ*M. Right, serial dilution of MBP with 10% PEG8000, condensates started appearing around 10 *µ*M, scale bar = 10 *µ*m.

To further investigate the effects of PEG on phase separation, we used FRAP on the droplets pre-formed with HP1α-KCK-Atto488 and DNA in the presence of 10% PEG·8000. In the presence of PEG·8000, only ∼56% of the bleached fluorescence recovered over the timescale of the experiment in contrast to ∼99% recovery observed in the absence of PEG·8000 (Figure 4C vs. 1H and Table 3). FRAP experiments on droplets formed with HP1α-KCK-Atto488 alone in the presence of PEG·8000 showed an even smaller fraction of fluorescence recovery (Figure 4C). These results suggest that 10% PEG·8000 substantially changes the dynamics of the HP1α proteins within the phase-separated droplet (Figure 4C vs. 1H and Table 3).

Given that all the constructs we tested contain IDRs, which often promote phase separation, we wanted to test the effects of PEG on a protein that lacks IDRs. We therefore chose Maltose Binding Protein (MBP), which is a canonical globular protein, often used to solubilize other proteins. Interestingly, we observed phase separation of MBP in the presence of PEG, starting at 10 *µ*M MBP (∼ 3-fold higher than HP1α and HP1α-16aa-mEGFP, but ∼2-fold lower than HP1α CSDm) (Figure 4E, Tables 1 and 2).

**Table 1.** Saturation concentration (*µ*M) of different proteins with and without crowding agents PEG8000 and PEG4000.

| Crowding Agent | Protein Alone |  |  |  |
| --- | --- | --- | --- | --- |
| | WT HP1α<br>(μM) | WT HP1α-<br>*I165E* (μM) | HP1α-16aa-mEGFP (μM) | MBP<br>(μM) |
| None | >400 | >150 | >500 | >80 |
| 10 % PEG 8000 | 3.3 | 22.5 | 3.3 | 10 |
| 10 % PEG 4000 | 3.3 | 30 | 1.1 |  |

**Table 2.** Saturation concentration (*µ*M) of different proteins plus DNA, with and without crowding agents PEG8000 and PEG4000.

| Crowding Agent | Protein + DNA |  |  |
| --- | --- | --- | --- |
| | WT HP1α<br>(μM) | WT HP1α-<br>*I165E*<br>(μM) | HP1α-16aa-MEGFP<br>(μM) |
| None | 3.3 | >150 | >450 |
| 10 % PEG 8000 | 3.3 | 5.6 | 3.3 |
| 10 % PEG 4000 | 3.3 | 10 |  |

### PEG·4000 leads to phase separation of protein, similar to the effects of PEG·8000

We also used PEG·4000 in addition to PEG·8000 to see if PEG at lower molecular weight has distinct effects on phase separation. We used 10% PEG·4000 throughout the assays to keep our comparisons under the same conditions. With 10% PEG·4000, untagged HP1α started phase separating at 3.3 *µ*M, with and without DNA, while HP1α-16aa-mEGFP alone has a saturation concentration of 1.1 *µ*M (Tables 1 and 2, Figure S2A). Similar to 10% PEG8000, 10% PEG·4000 is not sufficient to induce phase separation of DNA alone (Figure S2C). In the presence of 10% PEG· 4000, HP1α CSDm phase separates with and without DNA at 10 *µ*M and 30 *µ*M, respectively (Figure S2 D and E). Our findings suggest that both PEG4000 and 8000 induce phase separation of a variety of proteins irrespective of their ability to phase separate without PEG. Furthermore, under conditions containing PEG, HP1α proteins show a substantially reduced dependence on DNA for promoting phase separation.

## DISCUSSION

Understanding how the phase-separation behavior of proteins contributes to biological function requires an ability to visualize the dynamics of the corresponding proteins in cells. A common and powerful way of studying such dynamics relies on tagging the protein of interest with a genetically encoded fluorescent tag. However, addition of the tag has the potential to perturb the intrinsic phase-separation properties of the protein. Here we sought to identify a genetically encoded tagging arrangement for the heterochromatin protein HP1α that would minimally perturb its phase-separation properties *in vitro*. We found that both the linker length and the size of the tag matter for designing such a tagging arrangement. Through this study, we have identified one minimally perturbative tagging arrangement for HP1α, a C-terminal (holo) UnaG tag (13 kDa) with a 16 amino acid linker. We have further investigated the effects of PEG -based crowding agents to assess whether mimicking the molecular crowding in the nucleus can increase the repertoire of genetically encoded tags that allow HP1α phase-separation with comparable properties as untagged HP1α. Below we discuss the implications of both parts of our study for investigations of biological phase-separation.

In the context of C-terminally tagging HP1α, we find that the commonly used GFP class of tags interferes with phase-separation irrespective of whether we use an 8 or 16 amino-acid linker. We recognize that a more exhaustive search of linker lengths would be required to rule out mEGFP as an appropriate tag. However our finding that the HP1α-16aa-(holo)UnaG allows phase-separation with comparable properties as untagged or chemically tagged HP1α, suggests that the larger size of mEGFP compared to (holo)UnaG is responsible for the perturbative effects of the mEGFP-tagging arrangements. The HP1α-16aa-(holo)UnaG construct phase separates with a 2.7 kbp plasmid DNA, starting at ∼6.9 μM, which is within 3-fold of the saturation concentration observed with untagged HP1α (Figure 1B and G). Our FRAP studies indicate that HP1α-16aa-(holo) UnaG also has comparable dynamics to chemically tagged HP1α within the phases. While HP1α-16aa-(holo)UnaG shows a smaller fraction of fluorescence that recovers in the fast phase compared chemically tagged HP1α, overall the bleached spot recovers up to 94.07% within seconds (Table 3). While size is one difference between UnaG and mEGFP, it is possible that differences in other features between the two tags such as surface charge distribution and solubility also contribute to their differential effects on the phase-separation of HP1α.

**Table 3.** FRAP data of 5 conditions tested, HP1α-KCK-Atto488 with 2.7 kbp DNA, HP1α-16aa-(holo)UnaG with 2.7 kbp DNA, HP1α-KCK-Atto488 with PEG8000, and HP1α-KCK-Atto488 with 2.7 kbp DNA and PEG8000, respectively.

| | WT HP1α<br>KCK + DNA | HP1α-16aa-<br>holo UnaG | WT HP1α<br>KCK +<br>PEG8000 | WT HP1α<br>KCK +<br>DNA+<br>PEG8000 |
| --- | --- | --- | --- | --- |
| Fast Half Life<br>(sec) | 1.9 | 3.1 | 1.3 | 2.4 |
| Slow Half Life<br>(sec) | 55.9 | 21.1 | 300.9 | 278.5 |
| Percent fast (%) | 65.5 | 38.0 | 9.8 | 33.42 |
| Percent recovery<br>after plateau<br>reached (%) | 99 | 94.07 | 36.4 (Did<br>not plateau) | 56.6 |

Interestingly, the mEGFP tag did not interfere with DNA binding (Figure S1C and D) suggesting that the tag inhibits the higher order HP1α oligomerization that is essential for phase-separation. Additionally, we uncovered some differences in the effects of the apo and holo versions of the UnaG tag. Specifically, whereas HP1α-8aa-(apo)UnaG phase separates at a comparable saturation concentration as untagged HP1α, the HP1α-8aa-(holo)UnaG protein does not phase-separate up to 30 μM. While we do not understand the basis for this difference, we speculate that holo form of UnaG causes larger steric hindrance compared to the apo form and that extending the linker length to 16 amino acids relieves this steric hindrance. Consistent with this possibility, the secondary structure of holo UnaG shows that it is a well-folded beta-barrel protein, and it is possible that without the Br cofactor, UnaG is less well structured (15). This potential feature of disorder to order transitions in tags affecting phase-separation is also exemplified by our finding the AID tag promotes phase-separation of HP1α. This result is consistent with the patches of disorder predicted in the sequence of the AID. Binding of auxin presumably structures the AID tag. Overall, our results imply that understanding the biophysical effects of the tags in the context of the protein of interest is critical when choosing tags for studying phase-separation in cells. Each protein is expected to have a different tolerance for tags and possibly proteins with N- and C-terminal IDRs that participate in multi-valent interactions may be more intolerant of tags in the context of phase-separation.

For HP1α with large tags that cannot phase separate *in vitro* but can get recruited to heterochromatic sites, this is most likely due to three possibilities. First, if the tagged HP1α is exogenously over-expressed it may rely on endogenous untagged HP1α for phase separation. Second, interactions with other heterochromatin components may compensate for defects in the phase-separation of tagged HP1α. Third, the molecular crowding in the nucleus may facilitate phase separation and compensate for the inhibition introduced by the tag. Here we studied the third possibility by testing PEG8000 and PEG4000, reagents that are commonly used to mimic molecular crowdedness *in vitro*. We found that the presence of PEG promotes phase-separation of HP1α when tagged with proteins larger than UnaG. However, the presence of PEG also greatly diminished the contributions of two types of biologically relevant interactions. First, phase-separation of HP1α was no longer reliant on the presence of DNA. Second, mutating the CSD-CSD interface in HP1α, which inhibits phase-separation of HP1α and raises the saturation concentration by at least 40-fold, now raises the saturation concentration by only 6-fold. If PEG simply increased the local concentration of HP1α and DNA, then we would have expected the saturation concentrations for both the wild-type and CSDm HP1α to decrease, but the difference to remain at least 40-fold. The observation that the difference between the saturation concentrations of wild-type and CSDm HP1α is reduced to 6-fold suggests that the types of inter-molecular interactions with and without PEG are different. Consistent with this possibility, the dynamics of HP1α within PEG induced phases are different. Our FRAP studies indicate that a smaller population of bleached HP1α molecules recover in the presence of PEG. Prior work has shown that when part of a droplet formed from HP1α-KCK-Atto488 and DNA is subjected to photobleaching, the recovery of fluorescence occurs on the order of seconds without a substantial decrease in the total intensity of the droplet. This result implies that HP1α molecules can exchange on the order of seconds between droplets. However, we find in the presence of PEG the total intensity of the droplet decreases during recovery of the photobleached region. This difference suggests that in the presence of PEG, exchange of HP1α molecules between droplets is much slower than within a droplet.

Overall, our results suggest that PEG alters the protein-protein and protein-DNA networks within condensates and does not simply act as an inert crowding agent. Our findings are consistent with previous studies that have shown that PEG partitions within NPM1 and rRNA droplets and decreases the mobile fraction of protein within the condensates and more recent studies showing that PEG8000 can drive the phase-separation of a wide range of proteins (16, 18). Therefore, as suggested in these previous studies it is important to assess the biological relevance of PEG induced phase-separation with caution. The ambiguity of using PEG *in vitro* does not take away the importance of estimating how macromolecular crowding in cells affects saturation concentrations of proteins. In a cell, the added complexity of other interacting components is also likely to change the saturation concentration. Indeed, recent studies show that the saturation concentration of a fluorescently tagged N-Myc protein is ∼ 400 nM in mammalian cells but is at least 10-fold higher *in vitro* (27).

## MATERIALS AND METHODS

### Protein Purification

All HP1α tagged constructs with a 6x-His tag on the N-terminus were ordered from Genscript. Rosetta competent cells (Millipore Sigma 70954) were transformed with expression vectors for 6x-HIS HP1α, and BL21-Gold (DE3) competent cells (Aligent Technology 230132) were transformed with expression vectors for all other 6x-HIS HP1α tagged constructs (mEGFP, UnaG, AID-sfGFP, AID). Both cells were grown in 2xLB supplemented with 25 *µ*g/mL carbenicillin at 37°C to an OD600 of 0.3-0.4 then moved to 18°C and then allowedto grow till the OD600 reached 0.6-0.8, before inducing with 0.8 mM isopropy-*β*D-thiogalactopyranoside (IPTG). Cells were then grown overnight for 16-20 hours at 18°C, before pelleting at 4000xg for 30 mins. Cell pellets were collected using a spatula and flash frozen in liquid nitrogen and stored at -80°C until the day of purification. Each pellet was resuspended and thawed in lysis/wash buffer (1x PBS, 300 mM KCl,10% Glycerol, 7.5 mM Imidazole), supplemented with protease inhibitors (1 *µ*g/mL pepstatin A (Gold Biotechnology P-020-100), 1 mM phenylmethanesulfonyl fluoride (Sigma-Aldrich 78830), and 3 *µ*g/mL leupeptin (Sigma-Aldrich L2884)). Cells were lysed using a C3 Emulsiflex (ATA Scientific). Lysate was clarified by centrifugation at 30,000xg for 40 minutes in Oakridge tubes. The supernatant was bound to of Talon cobalt resin (Takara 635636) on rotation for 1-1.5 hrs. The bound resin was batch washed with the lysis/wash buffer for 60x the resin volume. Protein was eluted in of elution buffer (20 mM HEPES pH 7.5, 100 mM KCl, 400 mM Imidazole). TEV protease was added to the eluted protein and allowed to dialyze overnight in dialysis buffer (20 mM HEPES pH 7.5, 150 mM KCl, 3 mM DTT). The cleaved protein was filtered through a 0.22 *µ*m filter and then further purified through anion exchange, using a Mono Q 10/100 GL column (GE Healthcare). The protein was eluted using a salt gradient, from 80 mM to 1 M KCl over 20 column volumes in buffer containing 20 mM HEPES pH 7.5 and 3 mM DTT. The fractions containing protein were pooled and concentrated in a 10K Amicon Ultra-4 spin concentrator (Millipore 234753) down to 500 *µ*L. The protein was injected onto a Superdex-75 Increase 10/300 GL column (GE Healthcare 29148721) for size exclusion chromatography in SEC buffer (20 mM HEPES pH 7.5, 10% Glycerol, 300 mM KCl, 3 mM DTT). Fractions containing protein were pooled and concentrated again, aliquots were then flash frozen and stored at -80°C.

All HP1α constructs were purified as described above. Minor changes were made for the tagged constructs that are larger than 50 kDa. During concentration, a 30K Amicon Ultra-4 spin concentrator (Millipore 216718) was used instead of a 10K concentrator. For size exclusion chromatography, a Superdex-200 Increase 10/300 GL column (Cytiva 28-9909-44) was used instead of a Superdex-75 increase column.

### UnaG Labeling

A stoichiometry of 2:1 Bilirubin to protein was added to purified HP1α-8aa-UnaG and HP1α-16aa-UnaG. The protein-dye mixture was incubated for 15 minutes on ice before separating the unbound Bilirubin on a PD10 desalting column.

### Phase separation assays

Greiner Sensoplate™ glass bottom 384-well plates (Sigma-Aldrich M4187) were used for visualization of phase-separated droplets in this study. The wells in the plates were first washed 2 times with 100 *µ*L of water, then with 100 *µ*L of 2% Hellmanex (Sigma-Aldrich Z805939) for 30 minutes, followed by three water rinses. Next, 100 *µ*L of 0.5M NaOH was added to each well for 30 minutes, followed by another water rinse. Afterward, 70 *µ* L of 20 mg/mL PEG-silane MW-5000 (Laysan Bio MPEG-SIL-5000) dissolved in 95% EtOH was added to each well and allowed to sit overnight at 4°C, away from light. The next day, 30 *µ*L of 95% EtOH was added to each well and the wells were washed 2 times with 95% EtOH, but not left dry for more than 5 seconds. After the EtOH wash step, the wells were washed 3-times with water. Next, 100 *µ*L of 100 mg/mL of BSA (Sigma A4503-50G) was added to each well and allowed for a 30-minute incubation. The wells were washed extensively (5 times) with water and 2 times with assay buffer (20 mM HEPES pH 7.5, 70 mM KCl, 1 mM DTT).

Both HP1α constructs and DNA were dialyzed into phasing buffer (20 mM HEPES pH 7.5, 70 mM KCl, 1 mM DTT) overnight prior to experiment. Samples were calculated to be 2x each and added to PCR tubes on ice for 20 minutes for final concentration of 1x and a total volume of 15-30 *µ*L. For all condensates induced by PEG, 100 mg/mL was used as 10% PEG. Assay buffer remaining in the wells was rapidly removed, and samples were immediately added into the wells, one condition at a time. The droplets were allowed to settle at room temperature for 1 hour before imaging. All experiments were performed in triplicate. The settled non-fluorescent condensates were visualized by bright field microscopy at 20x magnification using the 6D/High throughput microscope at the Nikon imaging center at UCSF (except for MBP which was imaged at 40x magnification). Fluorescently tagged HP1α constructs were visualized using the same microscope but using the FITC channel instead of brightfield.

### FRAP assays and analysis

Droplets used for FRAP was prepared using the methods above, however, imaging was done using the Crest LFOV Spinning Disk/C2 Confocal microscope at 100x magnification at room temperature. For non-PEG condensates, a final concentration of 10 *µ*M of HP1α and HP1α-16aa-UnaG were mixed with 30 nM of 2.7 kbp DNA. For all PEG induced condensates, the same final concentration of HP1α, HP1α tag constructs, and DNA were used, with 10% PEG. All condensates were photobleached in the GFP-FRAP channel, using 10% 405 nm laser power. A round region of interest (ROI) was drawn on the area of interest for bleaching and assigned to the setting as described below. For all droplets, dwell time was set to 200 *µ* s. Three images were captured before photobleaching, for an interval of 300 msec, followed by 200 msec stimulation of bleaching. Without delay, 15 seconds of images were taken with 50 loops, and then 1 second intervals for more images for another 30 seconds. After stimulation, the same ROI was replicated two times, where one was placed in a background area, and the other one placed on a droplet that had not been bleached. All intensity values were measured using the Element software and exported to excel. To normalize intensity, an equation was used (Bleached ROI intensity – Background ROI intensity) / (non-bleached droplet ROI intensity - Background ROI intensity).

The data was averaged and fitted to the following equation:

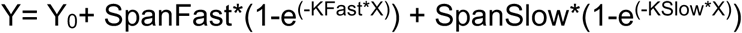

SpanFast=(Plateau-Y0)*PercentFast*.01

SpanSlow=(Plateau-Y0)*(100-PercentFast)*.01

Constraint: Plateau was set to 1

To collect Fraction long term recovery data, constraint was deleted and the same equations above were used.

### Electrophoretic mobility shift assays

Reactions contained HP1α or tagged HP1α at various concentrations (stated in figure legend), with 20 nM of Cy5 labeled 187 bp DNA in phasing buffer (20 mM HEPES pH 7.5, 70 mM KCl, 1 mM DTT). Reactions were carried out on ice but allowed to incubate at room temperature for 15 minutes before loading. A final concentration of 10% glycerol was added to reactions after incubation and bound and unbound DNA was separated by electrophoresis on a 6% acrylamide, 0.5X TBE native gel for 3 hours. The gel was imaged on an Amersham Typhoon laser-scanner platform (29187191) in the Cy5 channel. Fraction unbound was quantified in Fiji as a function of increasing concentration of protein and the following equation was fit to the data:

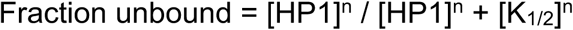

## ACKNOWLEDGEMENTS

We thank Upneet Kaur for helpful guidance on the writing of the manuscript and experimental design. We thank Emily Wong for experimental set up, CBX5 plasmid, and HP1α-KCK-Atto488. We thank Upneet Kaur and Elise Muñoz for the Cy5 labeled 187 bp DNA. We thank the Center for Advanced Light Microscopy (CALM) at UCSF for training on microscopes and troubleshooting. We thank Dr. Xiaokun Shu for suggesting the use of the UnaG tag and feedback on the manuscript. We thank Bo Huang and Kibeom Hong for helpful guidance on the FRAP experiments and feedback on the manuscript. We thank Emily Wong and Camille Moore for helpful feedback on the manuscript, and the Narlikar lab for stimulating discussions during the process of this study. This research was funded by grants from the NIH to G.J.N (U01DK127421 and R35GM127020).

