## Supplemental figures for "Understanding how genetically encoded tags affect phase separation by Heterochromatin Protein HP1α"

### SUPPLEMENTAL FIGURE LEGENDS

**Supplemental Figure 1.** Understanding how tags affect HP1α's biochemical properties (A and B) Bright field images of a concentration series of HP1α-8aa-Apo UnaG and HP1α-16aa-Apo UnaG with 30 nM 2.7kbp plasmid DNA, respectively. Scale bar = 20 $\mu$m. Red box indicates saturation concentration. (C and D) EMSA analysis of both wt HP1α and HP1α-16aa-mEGFP at 0, 0.4, 0.8, 1, 1.2, 1.4, 1.6, 2, 2.4, 2.8, 3.2, and 6.4 $\mu$M pre-incubated with 20 nM Cy5 labeled 187 bp DNA. (E and F) Quantified EMSA data of both wt HP1α and HP1α-16aa-mEGFP at varied concentrations bound to 20 nM Cy5 labeled 187 bp DNA.

**Supplemental figure 2.** PEG4000 induces phase separation of tagged and untagged HP1α. (A) From top to bottom, concentration series of WT HP1α and HP1α-16aa-mEGFP with 10% PEG 4000, respectively. (B) 150 $\mu$M of HP1α CSDm with 30 nM 2.7kbp DNA, no PEG. (C) 10% PEG4000 with 30 nM 2.7 kbp DNA. (D) A concentration series of HP1α CSDm with 10% PEG4000. (E) A concentration series of HP1α CSDm with 30 nM 2.7 kbp DNA and 10% PEG 4000. Scale bar = 20 $\mu$m. Red box indicates saturation concentration.

**Figure S1**

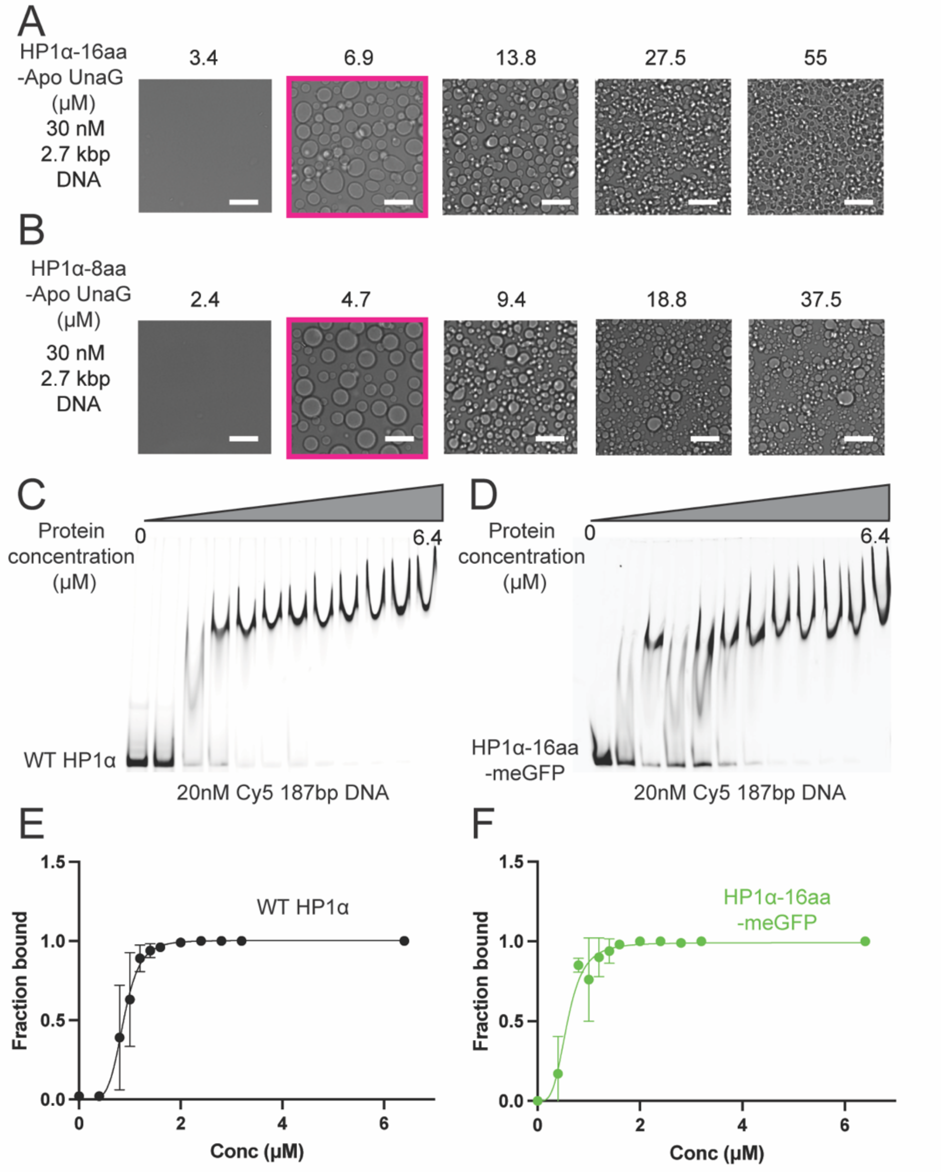
A
HP1α-16aa
-Apo UnaG
(μM)
30 nM
2.7 kbp
DNA
3.4
6.9
13.8
27.5
55
B
HP1α-8aa
-Apo UnaG
(μM)
30 nM
2.7 kbp
DNA
2.4
4.7
9.4
18.8
37.5
C
Protein
concentration
(μM)
0
6.4
WT HP1α
20nM Cy5 187bp DNA
D
Protein
concentration
(μM)
0
6.4
HP1α-16aa
-meGFP
20nM Cy5 187bp DNA
E
Fraction bound
Conc (μM)
WT HP1α
F
Fraction bound
Conc (μM)
HP1α-16aa
-meGFP

**Figure S2**

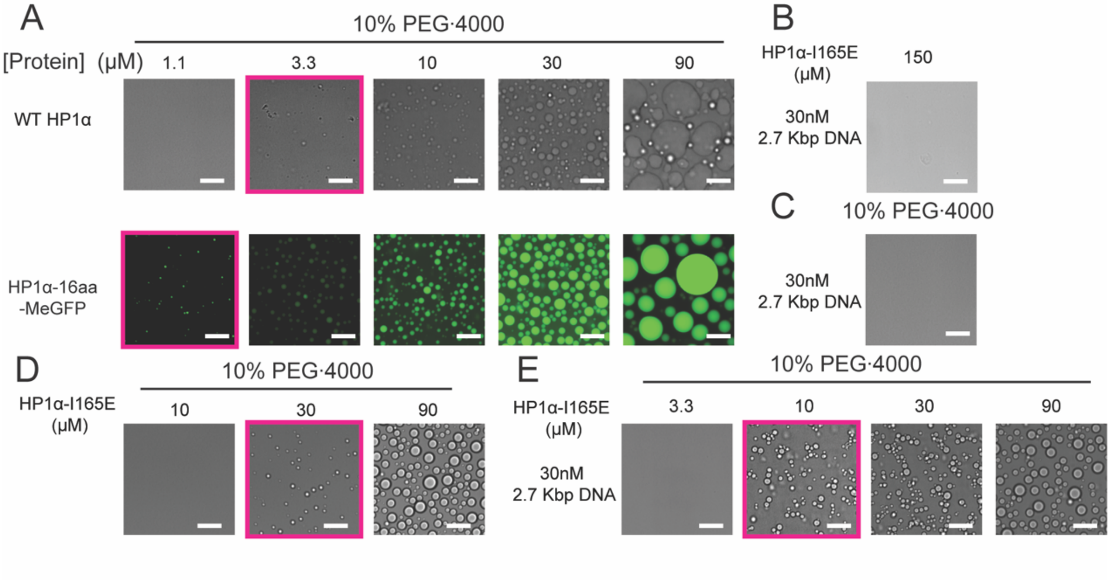

A
10% PEG·4000
[Protein] (μM) 1.1 3.3 10 30 90
WT HP1α
HP1α-16aa -MeGFP
B
HP1α-I165E (μM)
150
30nM 2.7 Kbp DNA
C
10% PEG·4000
30nM 2.7 Kbp DNA
D
10% PEG·4000
HP1α-I165E (μM)
10 30 90
E
10% PEG·4000
HP1α-I165E (μM)
3.3 10 30 90
30nM 2.7 Kbp DNA
